# The administration of high-mobility group box 1 fragment prevents deterioration of cardiac performance by enhancement of bone marrow mesenchymal stem cell homing in the delta-sarcoglycan-deficient hamster

**DOI:** 10.1101/390864

**Authors:** Takashi Kido, Shigeru Miyagawa, Takasumi Goto, Katsuto Tamai, Takayoshi Ueno, Koichi Toda, Toru Kuratani, Yoshiki Sawa

**Affiliations:** Department of Cardiovascular Surgery, Osaka University Graduate School of Medicine; Department of Stem Cell Therapy Science, Osaka University Graduate School of Medicine

**Author notes:** These authors contributed equally to this work.

## Abstract

**Objectives:** We hypothesized that systemic administration of high-mobility group box 1 fragment attenuates the progression of myocardial fibrosis and cardiac dysfunction in a hamster model of dilated cardiomyopathy by recruiting bone marrow mesenchymal stem cells thus causing enhancement of a self-regeneration system.

**Methods:** Twenty-week-old J2N-k hamsters, which are δ-sarcoglycan-deficient, were treated with systemic injection of high-mobility group box 1 fragment (HMGB1, n=15) or phosphate buffered saline (control, n=11). Echocardiography for left ventricular function, cardiac histology, and molecular biology were analyzed. The life-prolonging effect was assessed separately using the HMGB1 and control groups, in addition to a monthly HMGB1 group which received monthly systemic injections of high-mobility group box 1 fragment, 3 times (HMGB1, n=11, control, n=9, monthly HMGB1, n=9).

**Results:** The HMGB1 group showed improved left ventricular ejection fraction, reduced myocardial fibrosis, and increased capillary density. The number of platelet-derived growth factor receptor-alpha and CD106 positive mesenchymal stem cells detected in the myocardium was significantly increased, and intra-myocardial expression of tumor necrosis factor α stimulating gene 6, hepatic growth factor, and vascular endothelial growth factor were significantly upregulated after high-mobility group box 1 fragment administration. Improved survival was observed in the monthly HMGB1 group compared with the control group.

**Conclusions:** Systemic high-mobility group box 1 fragment administration attenuates the progression of left ventricular remodeling in a hamster model of dilated cardiomyopathy by enhanced homing of bone marrow mesenchymal stem cells into damaged myocardium, suggesting that high-mobility group box 1 fragment could be a new treatment for dilated cardiomyopathy.

## Introduction

Dilated cardiomyopathy (DCM) is one of the most common causes of heart failure and is associated with left ventricular dilatation and contractile dysfunction [1]. While significant improvements have been made in medical therapies, such as angiotensin-converting enzyme inhibitors and beta-blockers [2], and interventions, such as implantable cardioverter defibrillators [3] and cardiac resynchronization therapy [4], the prognosis for heart failure patients is still poor with 1-year mortality of 25–30% and a 50% survival rate at 5 years [5]. DCM remains the most common indication for cardiac transplantation but donor shortages have become a serious issue. To deal with this problem, several novel approaches using cell therapy have been developed in DCM patients with encouraging results [6–8].

Stem cells are an endogenous physiological healing mechanism of the body. A number of reports have suggested that damaged tissues may release various cytokines, which facilitate not only the mobilization of bone marrow-derived mesenchymal stem cells (BMMSCs) into the peripheral blood, but also their homing to sites of wound healing [9–11]. The enhancement of such healing mechanisms by drug administration might have beneficial effects in various diseases.

High-mobility group box 1 (HMGB1) is a non-histone nuclear protein that regulates chromatin structure remodeling by acting as a molecular chaperone in the chromatin DNA-protein complex [12]. Previous reports have demonstrated that endogenous platelet-derived growth factor receptor-alpha positive (PDGFRα^+^) BMMSCs accumulate in damaged tissue and contribute to regeneration in response to elevated HMGB1 levels in serum [13]. Moreover, systemic administration of HMGB1 further induces the accumulation of PDGFRα^+^ BMMSCs in the damaged tissue through CXCR4 upregulation, which is followed by significant inflammatory suppression [14].

Since BMMSCs have been reported to have therapeutic effect in DCM through paracrine effects [6,7], the above-mentioned “drug-induced endogenous regenerative therapy” might have effectiveness for DCM without supply of viable ex vivo cells. Recently, we developed a HMGB1 fragment containing the mesenchymal stem cell mobilization domain from human HMGB1. We hypothesize that systemic administration of this HMGB1 fragment attenuates the progression of myocardial fibrosis and cardiac dysfunction in a hamster model of DCM by recruitment of BMMSCs, promoting self-regeneration.

## Material and Methods

Animal procedures were carried out under the approval of the Institutional Ethics Committee (reference number 28-011-002). The investigation conformed to the “Principles of Laboratory Animal Care” formulated by the National Society for Medical Research and the “Guide for the Care and Use of Laboratory Animals” (National Institutes of Health Publication). All experimental procedures and evaluations were performed in a blinded manner.

### Experimental Animals

Male J2N-k hamsters, which are deficient in δ-sarcoglycan, were used for this study. J2N-k hamsters are an established animal model of DCM. They exhibit progressive myocardial fibrosis and moderate cardiac dysfunction at 8–9 weeks of age. Accordingly, the average life span of J2N-k hamsters is much shorter (approximately 42 weeks) than control hamsters (approximately 112 weeks) [15,16].

### HMGB1 Fragment

Mesenchymal stem cell mobilization domain from human HMGB1 was produced as “HMGB1 fragment” by solid-phase synthesis and provided by StemRIM (StemRIM Inc., Osaka, Japan).

The HMGB1 fragment was dissolved in phosphate buffered saline (PBS) to a concentration of 1 mg/ml before administration.

### Procedure of HMGB1 Fragment Administration

Male 19-week-old J2N-k hamsters were purchased from Japan SLC (Shizuoka, Japan). HMGB1 fragment (3mg/kg/day; HMGB1, n= 15) or PBS (3ml/kg/day; control, n=11) was administered for 4 consecutive days at the age of 20 weeks in the following manner: The external jugular vein was exposed by a median neck skin incision under 1.5% isoflurane anesthesia. Subsequently, HMGB1 fragment or PBS was injected through the external jugular vein. After the complete hemostasis, the skin incision was closed, and the hamsters were housed in a temperature-controlled cage.

### Transthoracic Echocardiography

Transthoracic echocardiography was performed to assess cardiac function using M-mode echocardiography with Vivid I (GE Healthcare) under isoflurane inhalation (1%). Diastolic and systolic dimensions of the left ventricle (LVDd/Ds), and left ventricular ejection fraction (LVEF) were measured before treatment, and reassessed at 4 and 6 weeks after treatment.

### Histological Analysis

The heart was excised under isoflurane anesthesia (5%) 6 weeks after treatment to perform histological and molecular biological analysis. The excised heart was fixed with either 10% buffered formalin for paraffin sections or 4% paraformaldehyde for frozen sections. The paraffin sections were stained with picrosirius red to assess the degree of myocardial fibrosis. The paraffin sections were used for immunohistochemistry and labeled using polyclonal CD31 antibody (1:50 CD31, Abcam, Cambridge, UK), anti-α-sarcoglycan (clone: Ad1/20A6; Novocastra, Weltzar, Germany), and anti-α-dystroglycan (clone: VIA4-1; Upstate Biotechnology, Lake Placid, NY) to assess capillary vascular density and the organization of cytoskeletal proteins. The paraffin sections were also labeled using rabbit monoclonal anti-CD106 antibody (ab134047, Abcam, Cambridge, MA) and goat polyclonal anti-PDGFRα (R&D). PDGFRα and CD106 are known to be expressed in BMMSCs and are commonly used as markers for mesenchymal stem cells (MSCs) [17,18]. The frozen sections were also used for immunohistochemistry and labeled with rabbit polyclonal anti-SDF1 antibody (ab9797, Abcam, Cambridge, MA) and mouse monoclonal CXCR4 antibody (4G10, sc-53534, Santa Cruz Biotechnology). The frozen sections were also stained with 4-hydroxynonenal to estimate lipid peroxidation [19], and dihydroethidium to estimate superoxide production [20].

More than 5 sections were prepared per specimen and 3 low power fields per section were analyzed and averaged. The fibrotic area, the expression of α-sarcoglycan and α-dystroglycan, and the 4-hydroxynonenal positive area were measured using Metamorph image analysis software (Molecular Devices, Inc., Downingtown, PA). The capillary density, the number of PDGFRα^+^ and CD106 positive (CD106^+^) cells, the number of CXCR4 positive (CXCR4^+^) cells and SDF-1 positive area (mm^2^), and the number of dihydroethidium positive dots were measured using the BZ-analysis software (Keyence, Tokyo, Japan).

### Transmission electron microscopy

Cardiac tissue was pre-fixed with Karnovsky fixative containing 2.5% glutaraldehyde, 2% paraformaldehyde in a 0.1 M (pH 7.4) cacodylate buffer for 2 hours at 4°C and post-fixed with 2% osmium tetroxide for 2 hours at 4°C. The samples were then immersed in 0.5% uranyl acetate for 3 hours at room temperature, dehydrated in ethanol (50%, 70%, 80%, 90%, 95%, and 100%) and propylene oxide, and embedded in epoxy resin. Semithin sections (0.5μm) were stained with 0.1% toluidine blue solution and examined under a light microscope. Ultrathin sections were made with a Leica ultramicrotome. These sections were counterstained with uranyl acetate and lead citrate, before examination with a Hitachi H-7100 electron microscope at 75 kV.

### Real-Time Polymerase Chain Reaction

Total RNA was extracted from cardiac tissue and reverse transcribed using Omniscript reverse transcriptase (Quiagen, Hilden, Germany). The resulting cDNA was used for real-time polymerase chain reaction with the ABI PRISM 7700 system (Applied Biosystems) and Taqman Universal Master Mix (Applied Biosystems, Division of Life Technologies Corporation, Carlsbad, Calif) and using hamster-specific primers for tumor necrosis factor-α stimulating gene 6 (TSG-6), vascular endothelial growth factor (VEGF), and CXCR4. Each sample was analyzed in duplicate for each gene studied. Data were normalized to glyceraldehyde-3-phosphate dehydrogenase expression level. For relative expression analysis, the ddCT method was used, and a sample from a control hamster was used as reference. The real-time polymerase reaction was also conducted using Fast SYBR Green Master Mix and primers designed for hepatic growth factor and glyceraldehyde-3-phosphate dehydrogenase as shown in Table 1. For relative expression analysis, we prepared a 5-fold serial standard curve using a sample from a HMGB1 hamster as reference.

**Table 1.**
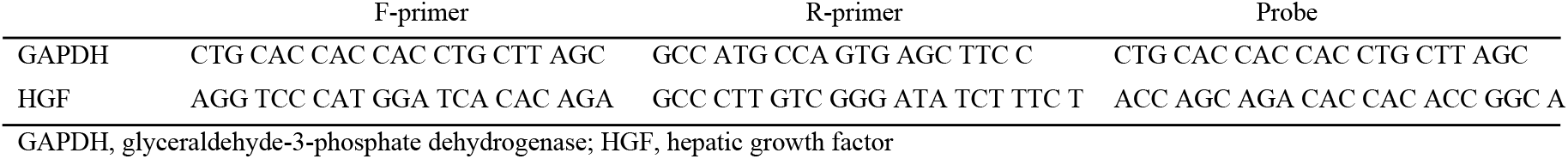
Forward and reverse primers and probe

### Evaluation of hamster prognosis after treatment

The life-prolonging effect of the HMGB1 fragment on J2N-k hamsters was assessed separately. Twenty-week-old J2N-k hamsters were treated with HMGB1 (HMGB1, n= 11) or PBS (control, n= 9) as described above. An additional treatment group received monthly administration of HMGB1 fragment 3 times, (monthly HMGB1, n= 9) to evaluate the long-term therapeutic effects of HMGB1 fragment (Fig 1). The animals were randomly allocated to each treatment group and housed after the initial treatment. The survival rate in the 3 groups was calculated by the Kaplan–Meier method using JMP Pro 12 (SAS Institute, Cary, NC, USA) and the significant difference between the groups was tested at 22 weeks (equal to 42 weeks of age, the average lifespan of J2N-k hamsters) by log-rank analysis.

**Fig 1.**
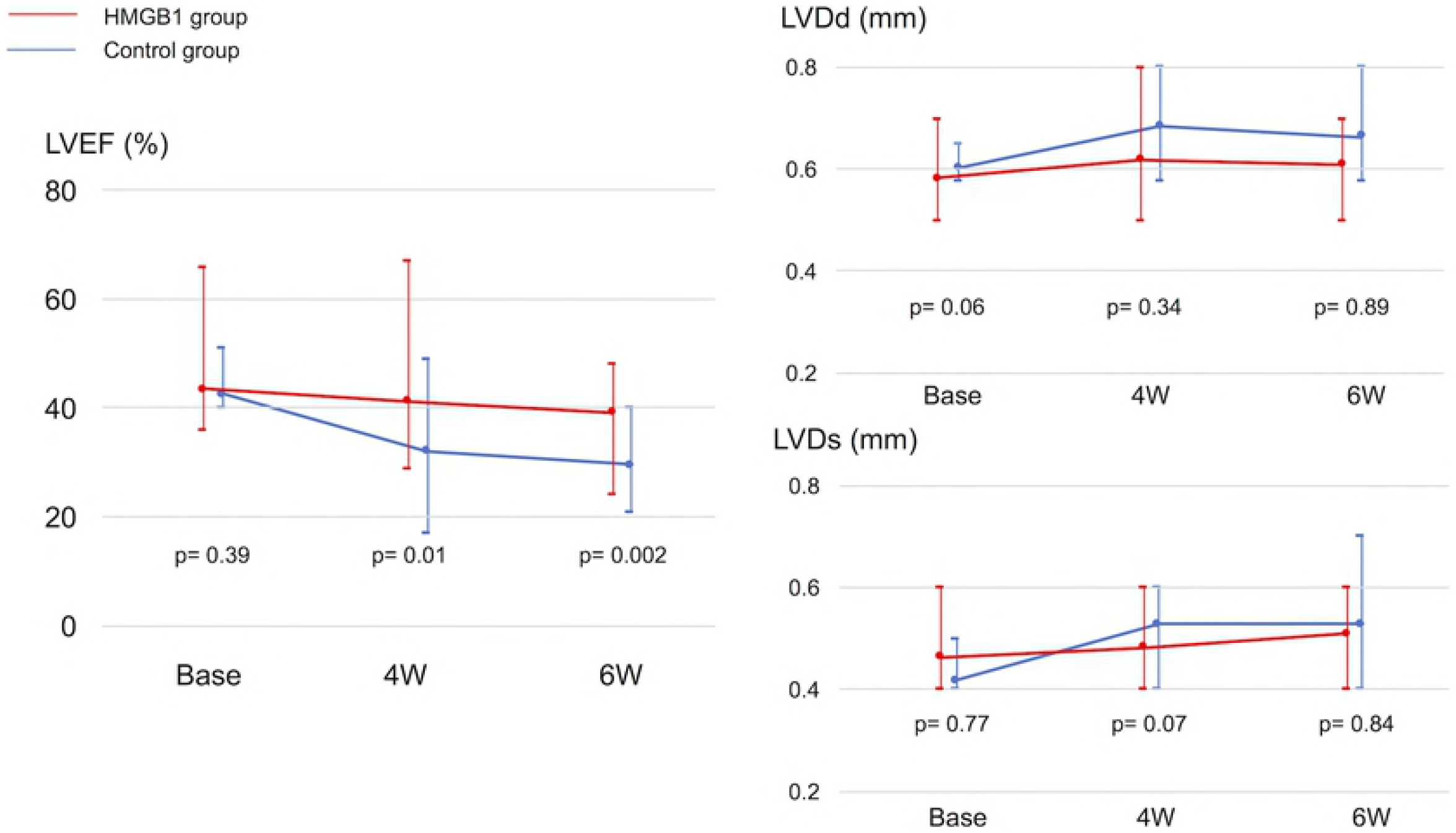
Changes in LVEF (a), LVDd (b), and LVDs (c) over time after the treatment. Diastolic and systolic dimensions of the left ventricle, and LVEF were measured before treatment, and reassessed at 4 and 6 weeks after treatment. The LVEF was significantly preserved until 6 weeks after the treatment in the HMGB1 group compared with the control group. LVEF, left ventricular ejection fraction; LVDd, left ventricular diastolic dimension; LVDs, left ventricular systolic dimension.

### Statistical Analysis

Continuous variables were summarized as means with standard deviations and compared using an unpaired t-test. Survival curves were prepared using the Kaplan–Meier method and compared using the log-rank test. All data were analyzed using JMP Pro 12. Differences were considered statistically significant at P-values < 0.05.

## Results

### Preserved Cardiac Performance with HMGB1 Fragment Administration

The functional effects of HMGB1 fragment on the DCM heart were assessed by transthoracic echocardiography over time. LVDd/Ds and LVEF at 20 weeks of age, just before the treatment, were not significantly different between the HMGB1 group and the control group. After treatment, echocardiography showed that the LVEF was significantly preserved until 6 weeks in the HMGB1 group compared with the control group (4 weeks: 43±8% vs 33±9%, p=0.01; 6 weeks: 41±7% vs 31±7%, p=0.0001, HMGB1 vs control, respectively) (Fig 1).

### Effect of HMGB1 Fragment on Myocardial Fibrosis

The degree of myocardial fibrosis 6 weeks after HMGB1 fragment treatment was assessed by picrosirius red staining and compared with control group. Quantification of fibrotic area confirmed that the degree of myocardial fibrosis was significantly reduced in the HMGB1 group compared with the control group (16.6±3.8% vs 22.7±5.4%, respectively, p=0.04) (Fig 2).

**Fig 2.**
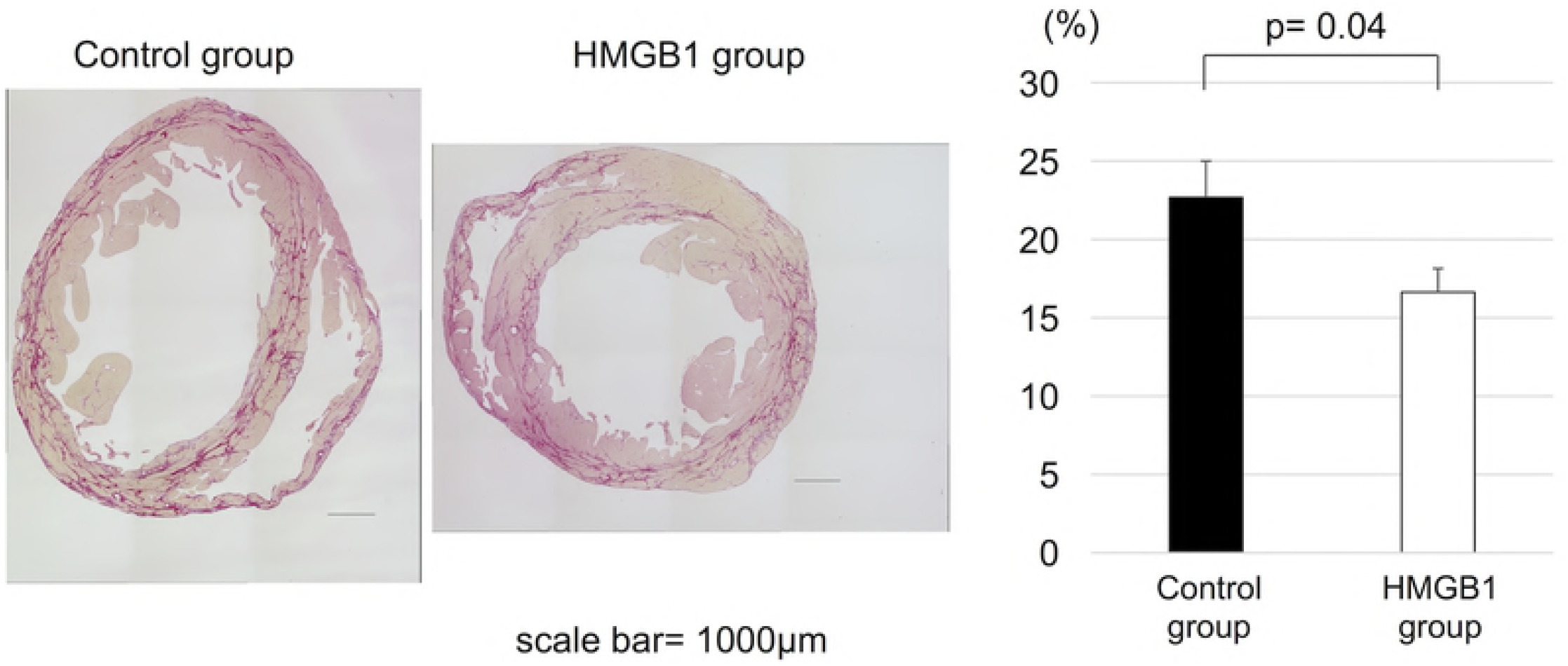
Suppression of myocardial fibrotic change in J2N-k hamsters by HMGB1 fragment. (a), Representative photomicrographs (×20, scale bar=1000μm) of picrosirius red staining. (b), Tissue sections were stained by picrosirius red and the fibrous area was quantified by image analysis. Percentage of myocardial fibrosis was significantly less in the HMGB1 group than in the control group. HMGB1, high-mobility group box 1.

### Increased Vasculature in the Heart After HMGB1 Fragment Administration

Capillary vascular densities 6 weeks after the treatment were assessed by CD31 immunostaining. In the HMGB1 group, the number of CD31 positive arterioles and capillaries was significantly increased compared with the control group (654±171 units/mm^2^ vs 484±74 units/mm^2^, respectively, p=0.02) (Fig 3).

**Fig 3.**
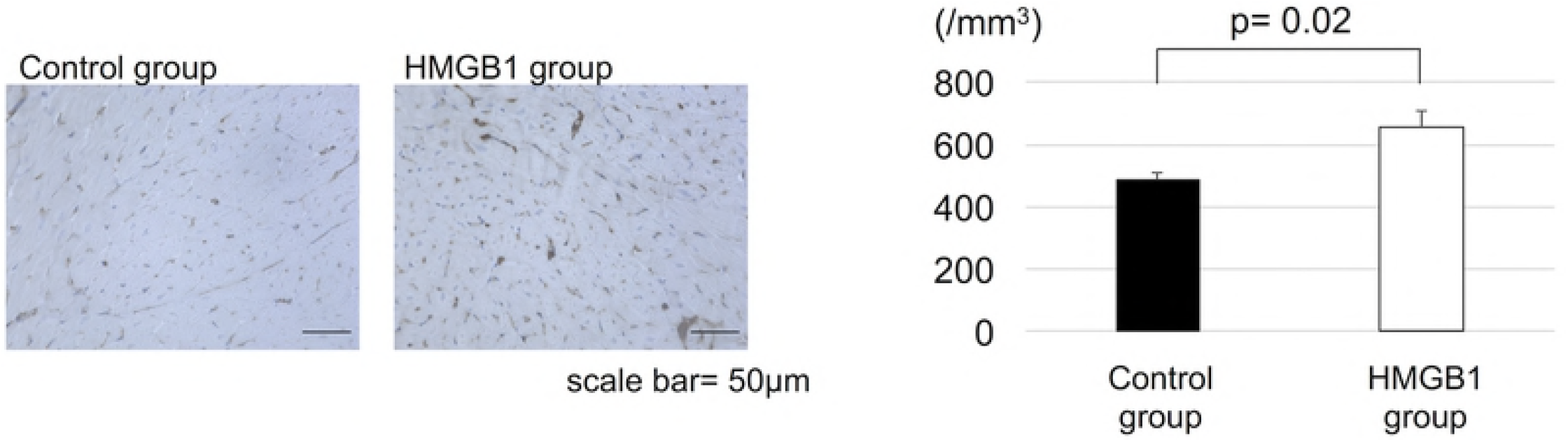
Increased myocardial capillary density in J2N-k hamsters by HMGB1 fragment. (a), Representative photomicrographs (×200, scale bar=50μm) of anti-CD31 staining. (b), Tissue sections were immunostained for CD31 and the capillary density was measured with the analysis software. The HMGB1 group showed significantly higher capillary vascular density than the control group. HMGB1, high-mobility group box 1.

### PDGFRα and CD106 Positive Cells in the Hearts

Immunohistochemistry showed that the number of PDGFRα^+^ and CD106^+^ cells in the heart tissue was significantly greater in HMGB1 group compared with the control group (12±5 cells /field vs 4±2 cells/field, respectively, p<0.001) (Fig 4).

**Fig 4.**
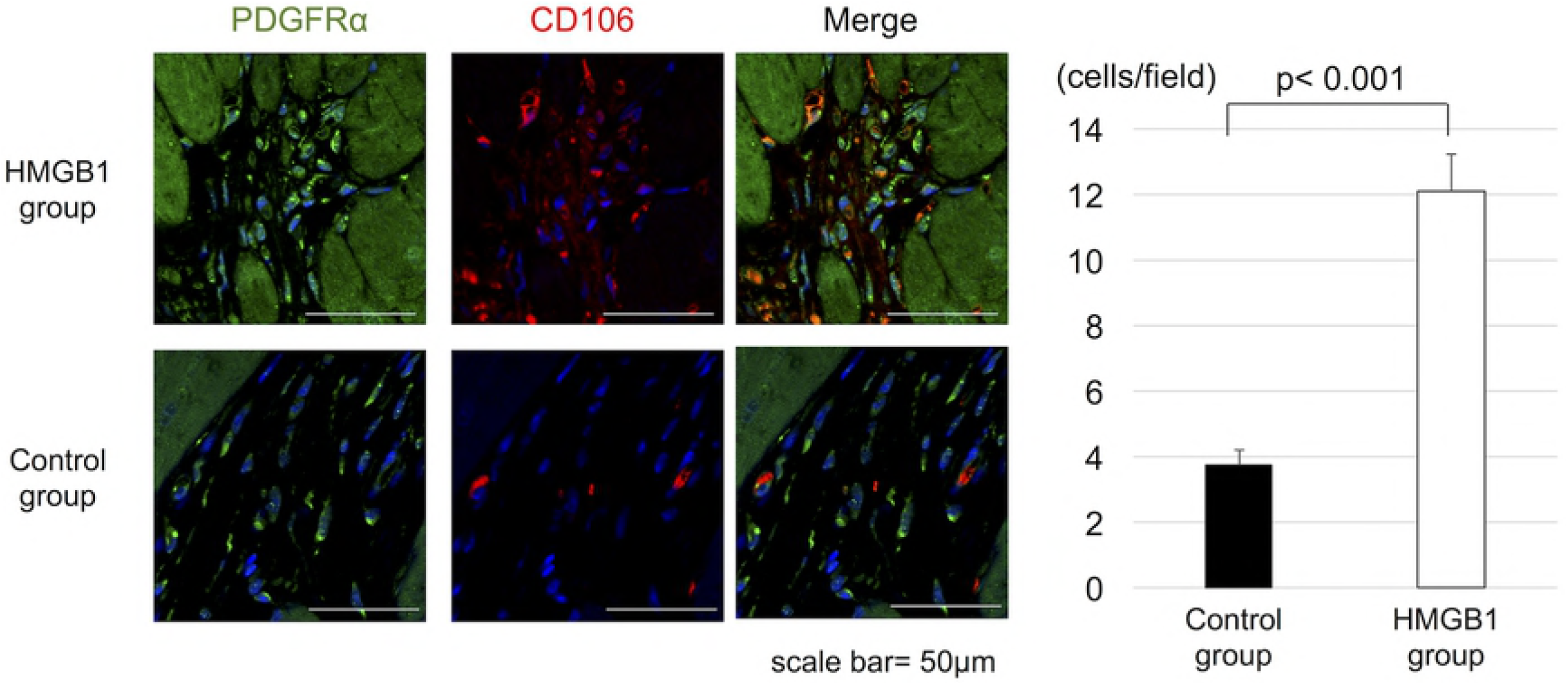
The increased accumulation of PDGFRα^+^ and CD106^+^ cells in the heart tissue by HMGB1 fragment. (a), Representative photomicrographs (×1000, scale bar=50 μm) of PDGFRα (green), CD106 (red) staining. (b), Tissue sections were immunostained for PDGFRα and CD106. The number of PDGFRα^+^ and CD106^+^ cells was measured with the analysis software. The HMGB1 group showed significantly increased numbers of PDGFRα^+^ and CD106^+^ cells than the control group. PDGFRα, platelet-derived growth factor receptor-alpha; HMGB1, high-mobility group box 1.

### Increased CXCR4 Positive Cells in the Heart after HMGB1 Fragment Administration

Immunohistochemistry showed significantly increased ratio of the number of CXCR4^+^ cells to SDF-1 positive area (mm^2^) in heart tissue in the HMGB1 group than in the control group (1.3±1.0 vs 0.3±0.1 cells/mm^2^, respectively, p=0.02) (Fig 5).

**Fig 5.**
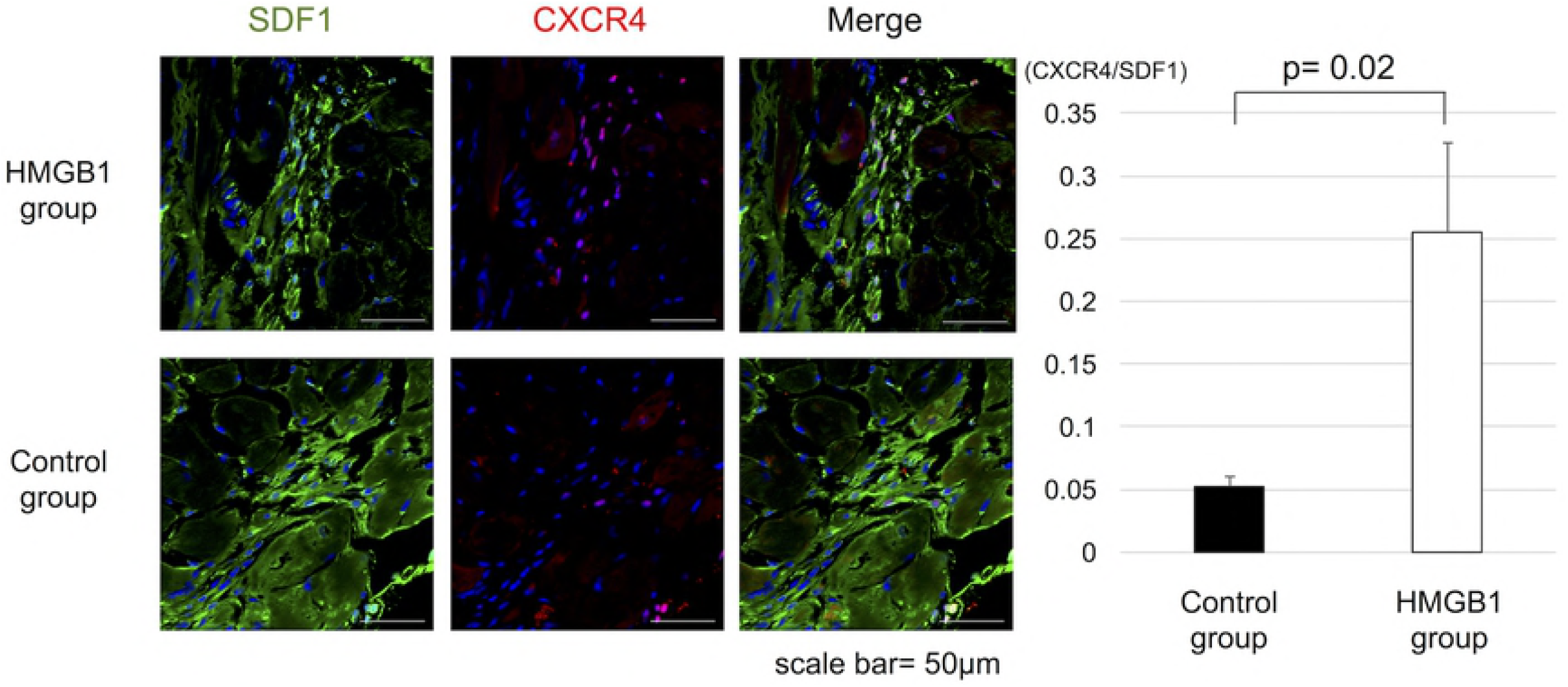
Increased CXCR4^+^ cells in SDF-1 positive area in the heart tissue by HMGB1 fragment. (a), Representative photomicrographs (×600, scale bar=50μm) of SDF-1 (green) and CXCR4 (red) staining. (b), Tissue sections were immunostained for CXCR4 and SDF-1. The number of CXCR4^+^ cells was measured with analysis software and SDF-1 positive area was measured with image analysis. The HMGB1 group showed significantly higher CXCR4^+^ cells to SDF-1 positive area (mm^2^) in heart tissue than the control group. HMGB1, high-mobility group box 1; SDF-1, stromal derived factor-1.

### Preservation of Cytoskeletal Proteins after HMGB1 Fragment Administration

Immunohistochemistry showed increased expression of α-sarcoglycan and α-dystroglycan in the basement membrane beneath the cardiomyocytes in HMGB1 group, whereas lower expression levels of these proteins was seen in the control group (α-sarcoglycan, 12.2±2.7% vs 2.8±1.4%, p<0.001, α-dystroglycan, 20.2±3.5% vs 8.3±1.8%, p<0.001, HMGB1 vs control, respectively) (Fig 6).

**Fig 6.**
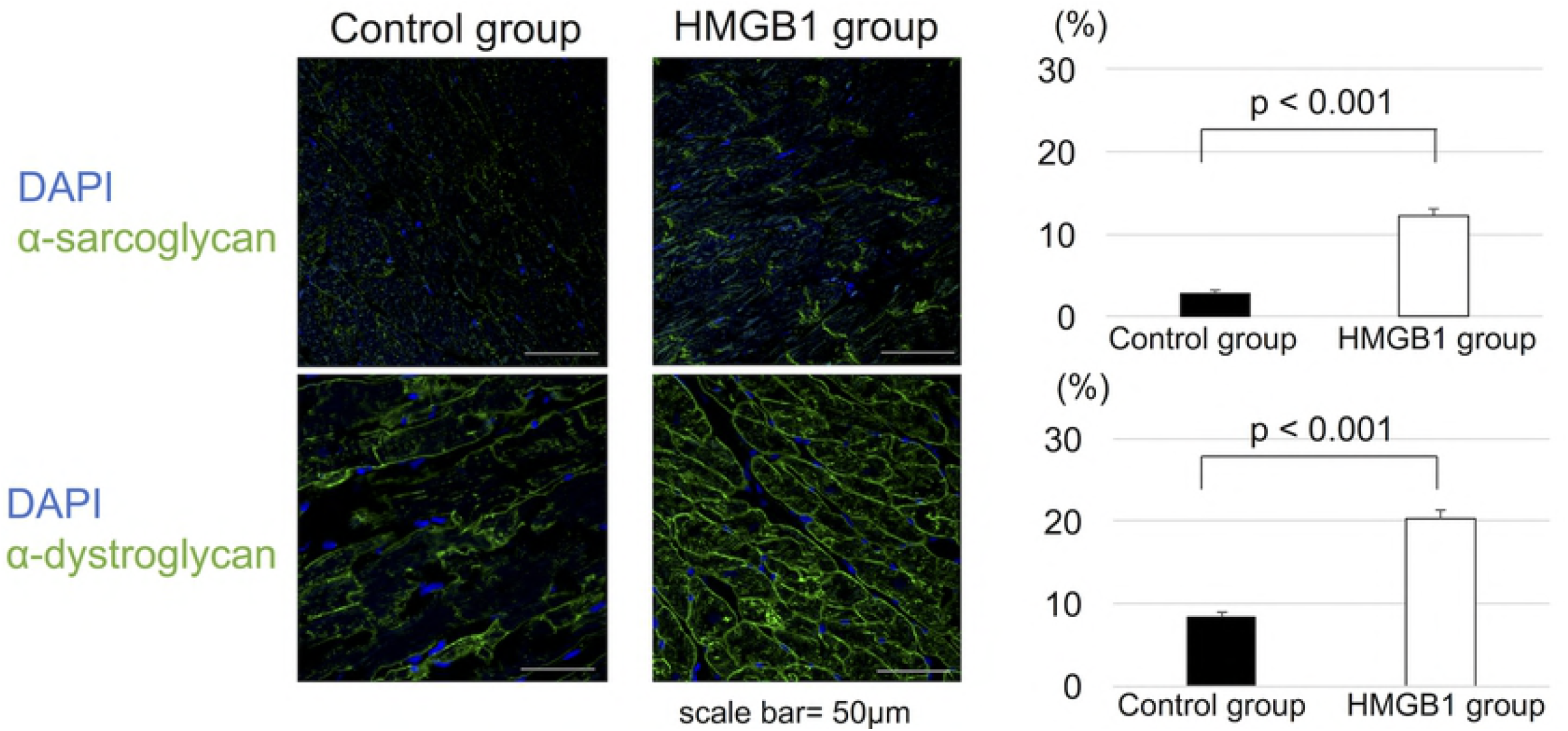
Immunostaining for alpha-sarcoglycan and alpha-dystroglycan in cardiomyocytes. (a), Representative photomicrographs (×600, scale bar=50μm) of immunostaining of α-sarcoglycan and α-dystroglycan in cardiomyocytes. (b), Quantitative analysis of immunohistologic signals showed significantly increased staining of both α-sarcoglycan and α-dystroglycan in HMGB1 group than the control group. HMGB1, high-mobility group box 1.

### Mitochondrial Ultramicrostructure

Transmission electron microscopy of the myocardium showed a relatively regular arrangement of mitochondrial cristae in the HMGB1 group. In contrast, the mitochondrial cristae were disordered in the control group (Fig 7).

**Fig 7.**
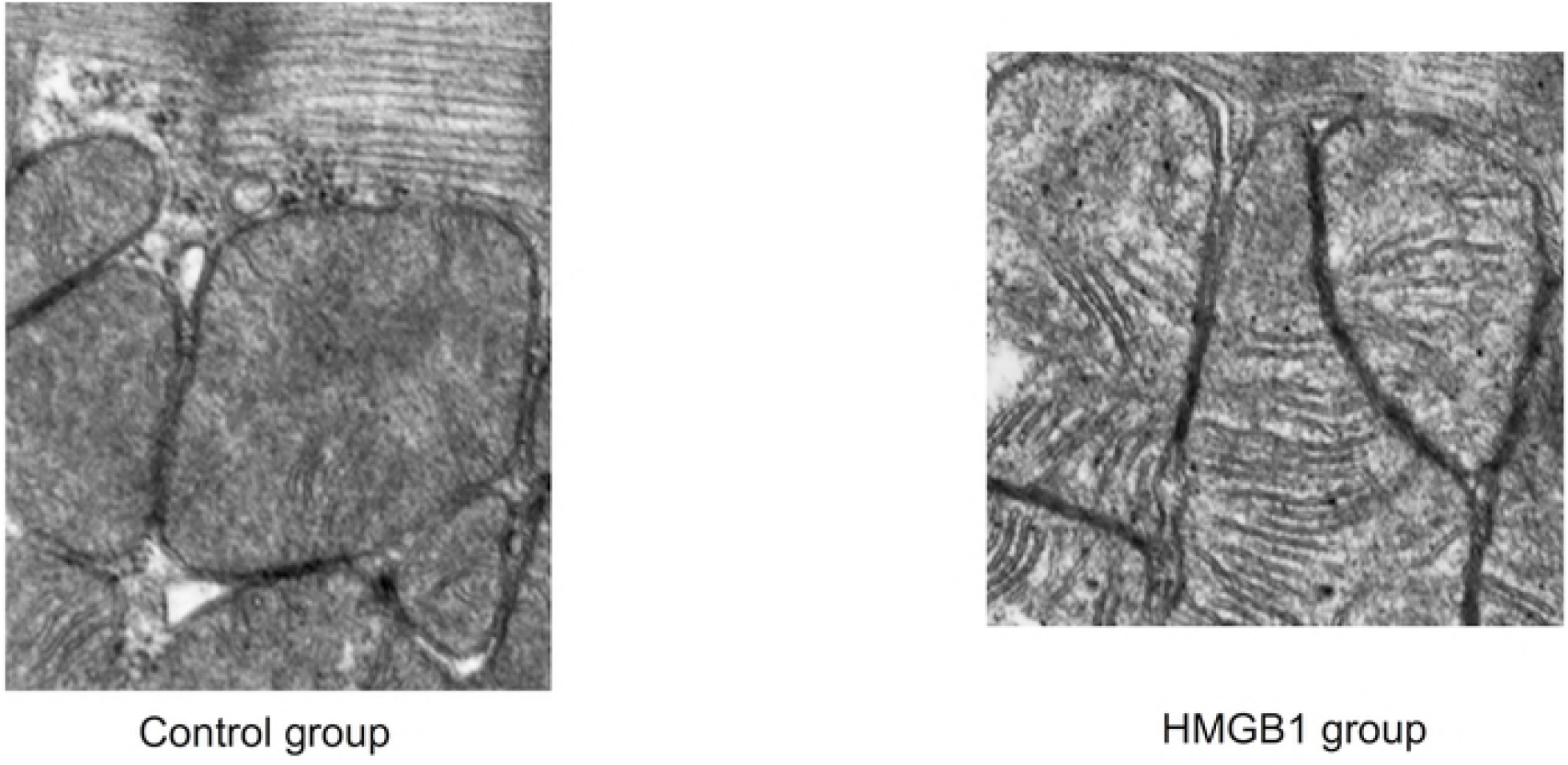
Mitochondrial ultramicrostructure was detected by TEM. Representative image (×12000) of mitochondrial morphology and cristae of myocardium in the HMGB1 group and the control group. The HMGB1 group showed relatively regular arrangement of mitochondrial cristae compared with the control group. TEM, Transmission electron microscopy; HMGB1, high-mobility group box 1.

### Effect of HMGB1 Fragment on Oxidative Stress in the Hearts

The lipid peroxidation and superoxide production were assessed by 4-hydroxynonenal staining and dihydroethidium staining, respectively. The results showed a trend towards reduced lipid peroxidation (3.5±2.4% vs 5.6±3.7%, p= 0.06, HMGB1 vs control) and a significant reduction in superoxide production in the HMGB1 group compared with control (219±32 units/mm^2^ vs 1185±97 units/mm^2^, respectively, p<0.0001) (Fig 8).

**Fig 8.**
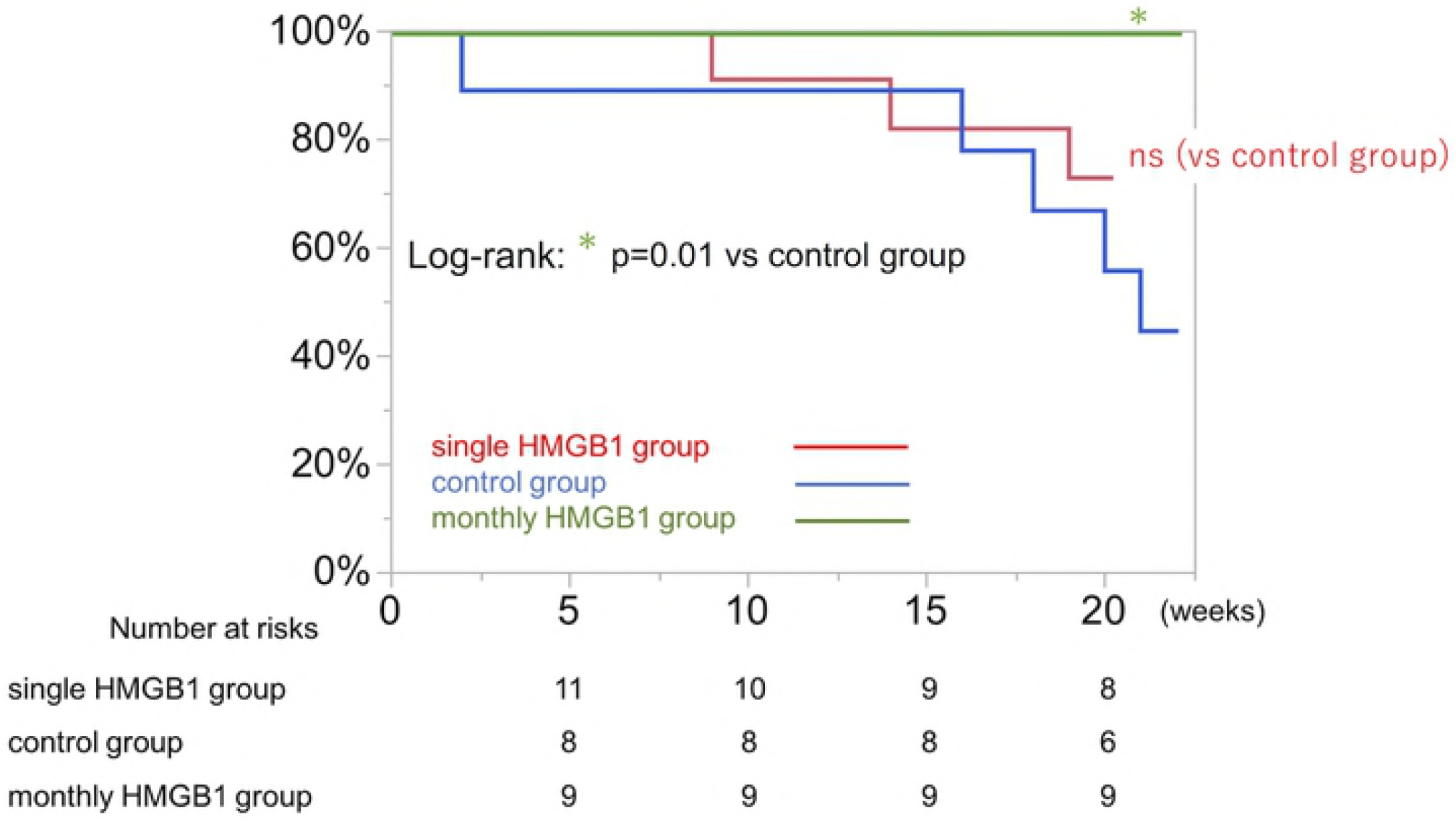
Decreased oxidative stress in the heart tissue by HMGB1 fragment. Representative photomicrographs of dihydroethidium staining (×400, scale bar=50μm) (a) and 4-hydroxynonenal staining (×100, scale bar=50μm) (b). Tissue sections were stained with dihydroethidium to estimate superoxide production, and 4-hydroxynonenal to estimate lipid peroxidation. The HMGB1 group showed significantly reduced production of superoxide (c) and a trend towards reduced lipid peroxidation (d) compared with the control group. HMGB1, high-mobility group box 1.

### Upregulated TSG-6, VEGF, HGF, and CXCR4 in the Heart after HMGB1 Fragment Administration

Real-time PCR was used to quantitatively assess the expression levels of BMMSC-derived factors, such as VEGF, TSG-6, HGF, and CXCR4. Intramyocardial mRNA levels of VEGF, TSG-6, and HGF were significantly upregulated in HMGB1 group compared with the control group (TSG-6, 1.5±0.6 vs 1.1±0.2, p= 0.03, VEGF, 1.3±0.4 vs 1.0±0.2, p= 0.04, HGF, 3.2±2.3 vs 1.3±0.6, p=0.02, HMGB1 vs control, respectively). The intramyocardial mRNA levels of CXCR4 in the HMGB1 group showed a trend towards increased expression compared with control (1.5±0.4 vs 1.2±0.3, respectively, p=0.06).

### Survival Benefit of Monthly HMGB1 Administration

Survival of J2N-k hamsters was assessed using the Kaplan–Meier method.

There was no significant difference in survival between HMGB1 and control. In contrast, the monthly HMGB1 group all survived to the full 42 weeks, and they showed significantly improved survival rate compared with control group (log-rank p=0.001) (Fig 9).

**Fig 9.**
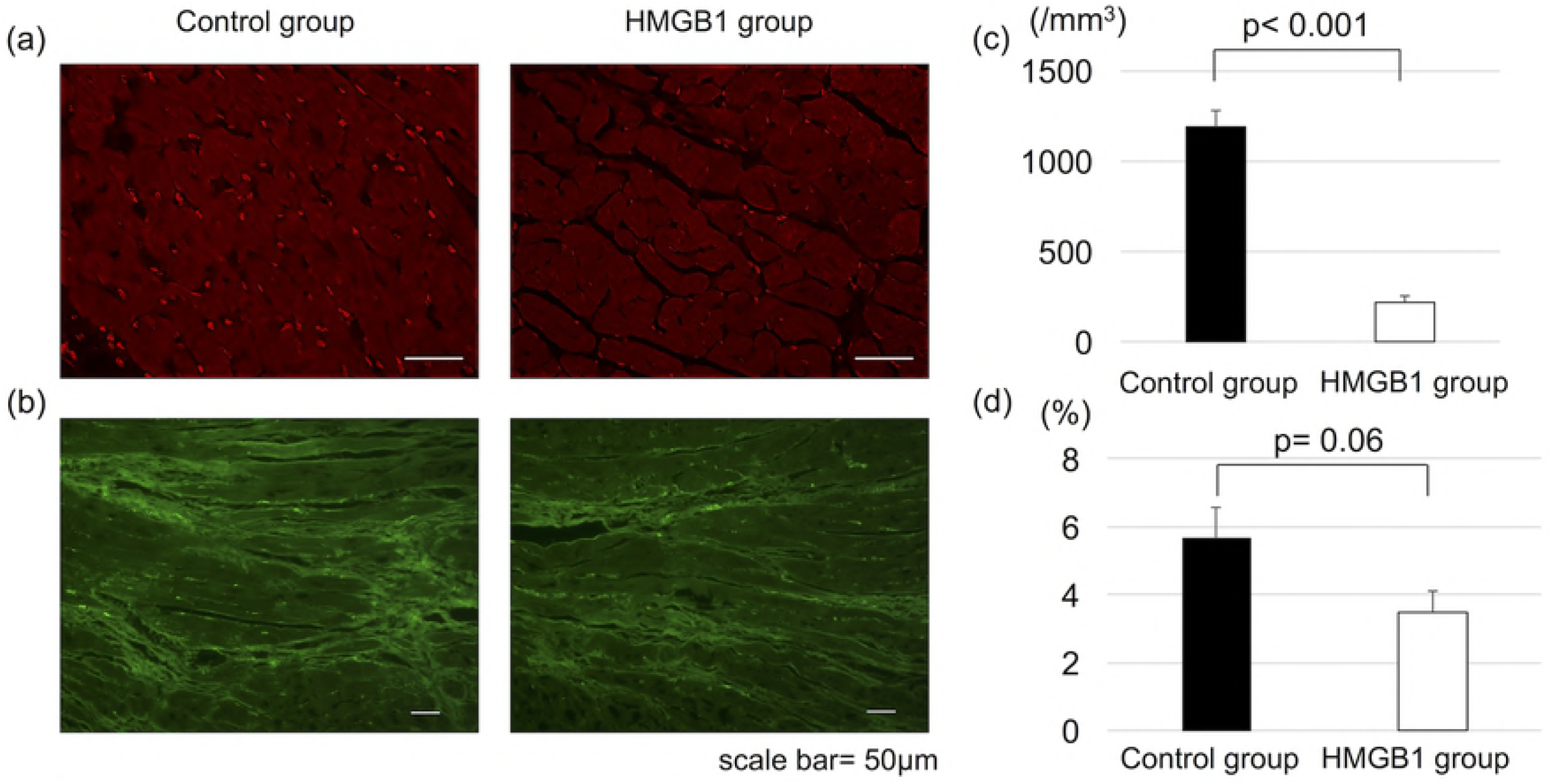
Survival after each treatment assessed by the Kaplan–Meier method. There was no significant difference between the single HMGB1 treatment (n=11) group and the control group (n=9), whereas the monthly HMGB1 group (n=9) showed a significantly greater survival rate than control (p=0.01).

## Discussion

In the present study we have shown that, first, systemic administration of HMGB1 fragment leads to the accumulation of PDGFRα^+^ and CD106^+^ cells in damaged myocardium possibly through the SDF-1/CXCR4 axis and upregulated expression of cardioprotective factors such as TSG-6, VEGF, and HGF in the heart tissue of J2N-k hamsters. Second, the myocardial histology in the HMGB1 group demonstrated significantly decreased fibrosis, increased capillary vascular density, decreased oxidative stress, and well-organized cytoskeletal proteins compared with the control group. Finally, cardiac function was significantly preserved after HMGB1 fragment administration and the survival benefit was shown with monthly HMGB1 fragment treatment.

The present study demonstrates the feasibility of “drug-induced endogenous regenerative therapy” using an HMGB1 fragment in a hamster model of DCM. While the precise mechanism remains unclear, it is well known that HMGB1 acts as a chemoattractant for MSCs [13,14,21]. Systemic HMGB1 administration has been reported to induce accumulation of PDGFRα^+^ cells around blood vessels in the bone marrow and significant increases in these cells in the peripheral blood [13]. In addition, the enhancement of CXCR4 expression with HMGB1 treatment promotes the local migration to damaged tissue through the SDF-1/CXCR4 axis, which might be essential in DCM [22] as well as ischemic cardiomyopathy [23–25].

While PDGFRα^+^ BMMSCs might be the predominant cell population mobilized by administration of HMGB1 fragment and therefore exerting therapeutic effects on damaged myocardium, it has been suggested that PDGFRα^+^ MSCs include other defined subpopulations with distinct functions [26]. As HMGB1 is also reported to induce other cell types [27], the accumulated cells in damaged heart tissue after HMGB1 administration might be highly heterogeneous and it will therefore be important to identify in the future, specific PDGFRα^+^ subpopulations induced by HMGB1 which have therapeutic benefits.

Paracrine signaling is a well-investigated mechanism of protective effects exhibited by BMMSCs on surrounding cells [28–32]. TSG-6 plays a key role in the anti-inflammatory effects of BMMSCs [31,33]. TSG-6 attenuates oxidative stress through activation of CD44 [34,35], and downregulates TGF-β by suppressing plasmin activity [33], which could result in decreased myocardial fibrosis. Since increased oxidative stress is one of the essential factors in the pathogenesis of myocardial fibrotic changes in J2N-k hamsters [16], our results suggest that HMGB1 fragment administration could become a substantial therapy for DCM. VEGF has been known to promote angiogenesis in ischemic conditions [29,36,37], which might have a beneficial effect on the defective vascularization within the left ventricle, which is associated with the pathophysiology of DCM [38,39]. HGF is known to be a putative paracrine mediator in cardiac repair mechanisms of BMMSCs [40]. Our group has previously reported that HGF induced the appropriate microenvironment for extracellular matrix remodeling, including strong expression of cytoskeletal proteins in damaged myocardium [41,42].

No significant difference in survival was observed between the HMGB1 and control groups, however, animals that received monthly HMGB1 treatment showed significantly better survival compared with control. The therapeutic benefits of HMGB1 fragment might be sustained by repeated administration in J2N-k hamsters and further investigation concerning the optimal dose and interval of administration of HMGB1 fragment will be needed for the clinical use of HMGB1 fragment in DCM patients.

## Conclusion

Systemic HMGB1 fragment administration attenuates the progression of left ventricular remodeling in a hamster model of DCM by enhanced homing of BMMSCs into damaged myocardium, suggesting that HMGB1 fragment could be beneficial in the treatment of DCM.

## Acknowledgments

We thank Hanne Gadeberg, PhD, from Edanz Group (www.edanzediting.com/ac) for editing a draft of this manuscript.

